# Anti-proliferative effect of potential LSD1/CoREST inhibitors based on molecular dynamics model for treatment of SH-SY5Y neuroblastoma cancer cell line

**DOI:** 10.1101/2022.05.23.493055

**Authors:** Hiba Zalloum, Waleed Zalloum, Tareq Hameduh, Husam ALSalamat, Malek Zihlif

## Abstract

Lysine-specific demethylase is a demethylase enzyme that can remove methyl groups from histones H3K4me1/2 and H3K9me1/2. It is expressed in many cancers, where it impedes differentiation and contributes to cancer cell proliferation, cell metastasis and is associated with inferior prognosis. LSD1 is associated with its corepressor protein CoREST, and utilizes tetrahydrofolate as a cofactor to accept CH2 from the demethylation process. The fact that the cofactor is best bound to the active site inspired us to explore its interactions to LSD1/CoREST enzyme complex utilizing molecular dynamics simulation, which aids designing novel and potent inhibitors. We have implemented a previously derived model from the MD simulation study and the key contacts to the active site in a subsequent structure based drug design and in-silico screening. In silico mining on National Cancer Institute (NCI) database identified 55 promising and structurally diverse inhibitors toward LSD1/CoREST complex. The anti-proliferative activities of the identified compounds were tested against neuroblastoma SH-SY5Y cancer cell line which known to highly express LSD1/CoREST complex. Applying the abovementioned molecular modeling procedure yielded Four compounds of LSD1/CoREST inhibiters with IC50 <2µM. The four lead compounds were tested against SH-SY5Y neuroblastoma cell line that known to express high level of LSD1 and illustrated a potent activity with an IC50 ranging from 0.195 to 1.52µM. To estimate the toxicity of the selective leads, they were tested against normal fibroblast cells and scored a relatively high IC50 ranging from 0.303 to ≥100µM. These compounds are excellent candidates treating cancers that overexpress the LSD1 enzyme.

## 1. Introduction

Epigenetics has emerged as the study of modifications of gene expression without altering the genetic material [1-4]. Epigenetic dysregulation contributes to the irregular gene expression programs of cancer, where it has an important mechanism in cancer initiation and progression [5, 6]. Mechanism that controls DNA and histone modifications has become a major focus for cancer targeted therapies [7-10].

Lysin specific histone demethylase (LSD1) was the first discovered histone demethylase in 2004, it plays an important role in normal and malignant cells [11, 12]. LSD1 is a flavin-dependent demethylase that belongs to the flavin adenine dinucleotide (FAD) family, it catalyzes the oxidative demethylation of mono -and di-methyl lysine residues of histones specifically at the H3K4 and H3K9 positions, as well as non-histone protein substrates, such as p53 [13, 14] DNMT1 [15], E2F1 [16], HIF-1 [17] and STAT3 [18], which results in transcriptional repression or activation [19-21]. Lysine-specific demethylase 1 (LSD1, also known as KDM1A and AOF2) is a chromatin-modifying activity that catalyzes the removal of methyl groups from lysine residues in histone and non-histone proteins, regulating gene transcription [22].

LSD1 is frequently found associated with other transcriptional factors such as its Co-Repressor for Element-1-Silencing Transcription factor (CoREST) to regulate variety of genes including the expression of tumour suppressor gene [3, 23]. Accordingly, LSD1/CoREST complex is considered as an important intracellular epigenetic target for the development of new anticancer drugs by reactivating the silenced tumor suppressor gene without destroying the gene itself. This would target cancerous rather than normal cells, which potentially enables selectively targeting cancer [14, 24, 25]. Abnormal overexpression of LSD1 was found in a variety of solid tumors, including neuroblastoma, breast, prostate, bladder, lung, liver, and colorectal tumors [20, 21, 26-29], where it inhibits differentiation, and enhances proliferation, invasiveness, and cell motility, and also worsens prognosis [30, 31]. Therefore, considerable interest emerged in the LSD1 inhibition as potential anti-cancer therapeutic strategy.

LSD1/CoREST complex uses tetrahydrofolate (THF) as a cofactor, where it accepts the methyl group resulted from the demethylation process of the methylated lysine residue [32, 33]. The fact that the cofactor is best bound to the active site inspired us to explore its interactions to LSD1/CoREST enzyme complex utilizing molecular dynamics simulation, which aids designing novel and potent inhibitors. Also, the conformational existence of the enzyme complex bound to the cofactor has been investigated [33]. In our initial work we have implemented molecular dynamics simulation to find a possible LSD1 protein conformation that has important role to inhibit this enzyme, and to determine the key contacts between the ligand and the active site of the enzyme. This followed by structure-based design and in silico screening revealed several new chemical entities with a potential inhibitory effect of LSD1

To date, all compounds that have been advanced into clinical trials are covalent-binding irreversible LSD1 inhibitors, with poor selectivity and toxic side effects. The design of highly potent and specific reversible LSD1 inhibitors for cancer therapy is still challenging and valuable.

In this study we minted to apply the mentioned molecular modeling procedure to identify the potential LSD1/CoREST inhibitors. Farther, we aimed to test the potency and the safety of such inhibitors against human neuroblastoma and fibroblast cells lines. The neuroblastoma was chosen because it does represent the most common cancer amongst those cancers that highly express LSD1 enzyme.

## 2. Material and Methods

### 2.1 Virtual Screening

The initial LSD1/CoREST enzyme structure used for the virtual screening was chosen to be the structures that were resulted from the MD simulation according to our previously published research [34]. The results of our previous research showed that the cofactor tetrahydrofolate binds at two different sites, 1 and 2. Accordingly, we used the RECEPTOR module of OpenEye Scientific Software Inc. to prepare the active sites of both structures for docking [35, 36].

The NCI database was used as input structures for the virtual screening, where the conformers of the NCI members were generated using Omega 2.5.1.4 module of the OpenEye Scientific Software Inc. using default parameters, except the search force field which was set to MMFF94 [35-37]. The tetrahydrofolate cofactor was docked into the sites 1 and 2 using ChemGauss4 scoring function to find the best docking parameters for the virtual screening [38-41]. The HYBRID module of OpenEye Scientific Software was used as docking engine. The default parameters were able to regenerate the tetrahydrofolate poses represented in the MD structures for both sites. Accordingly, we used the parameters of conformer generation and docking for the process of virtual screening using NCI database. Then, the highest score compounds, based on the ChemGauss4 scoring function, were selected and ordered from NCI for their experimental test as LSD1/CoREST inhibitors.

### 2.2 Cell Lines

SH-SY5Y neuroblastoma and dermal fibroblasts (<u>BJ ATCC® CRL-2522)</u> cell lines were grown and maintained in an incubator with a humidified atmosphere of 5% CO2 at 37°C. All were purchased from the American Type Culture Collection ATCC (Manassas, VA). Dermal fibroblasts were grown in Dulbecco’s modified Eagle’s medium (DMEM) with the same supplements added as with RPMI. SH-SY5Y cells were grown in a 1:1 mixture of ATCC-formulated Eagle’s Minimum Essential Medium and F12 Medium with 10% fetal bovine serum. All cell lines were grown in Corning® T-75 flasks where medium renewal was carried out every 2 -3 days and subculturing once every 4 days.

### 2.3 Antiproliferative assay

Both cell lines were washed with Phosphate Buffer Saline (PBS) and suspended for cell culturing using 0.25% trypsin 1X, 0.53 mM EDTA solution. Cells were then counted using Trypan blue and a haemocytometer and plated in the relevant medium at the appropriate seeding density into 96 well microtiter plates. The seeding densities for each cell line were determined by assuring that the cell cultures did not become confluent before carrying out the assay. After plating, the cells were incubated for 24 hrs to allow the cells to become attached to the bottom of the wells. They were then treated with and without drug and incubated with or without drug for 72 hrs. This allows time for cell proliferation and drug induced cell death to occur, as well as levels of enzymatic activity to drop which in turn generates formazan, the product from the MTT substrate. After 72 hrs, 100ul of media was removed from the wells and 15ul of yellow tetrazolium MTT (3-(4, 5-dimethylthiazolyl-2)-2,5-diphenyltetrazolium was added to each well. The plates were then incubated for 3 hrs. Once 3 hrs have passed, 100ul of solubilisation solution was added to each well to stop the reaction. The plates were then left for half an hour to allow the formazan crystals to dissolve, following which spectrophotometric absorbance was read at 540nm using an ELISA reader. The data of the MTT cell proliferation assay were manually analyzed using the GraphPad PRISM®8.0 software (GraphPad Software, Inc.). The inhibitory concentration (IC_50_) values, were calculated from the logarithmic trend line of the cytotoxicity graphs. **Figure 1** present the study flowchart.

**Fig 1:**
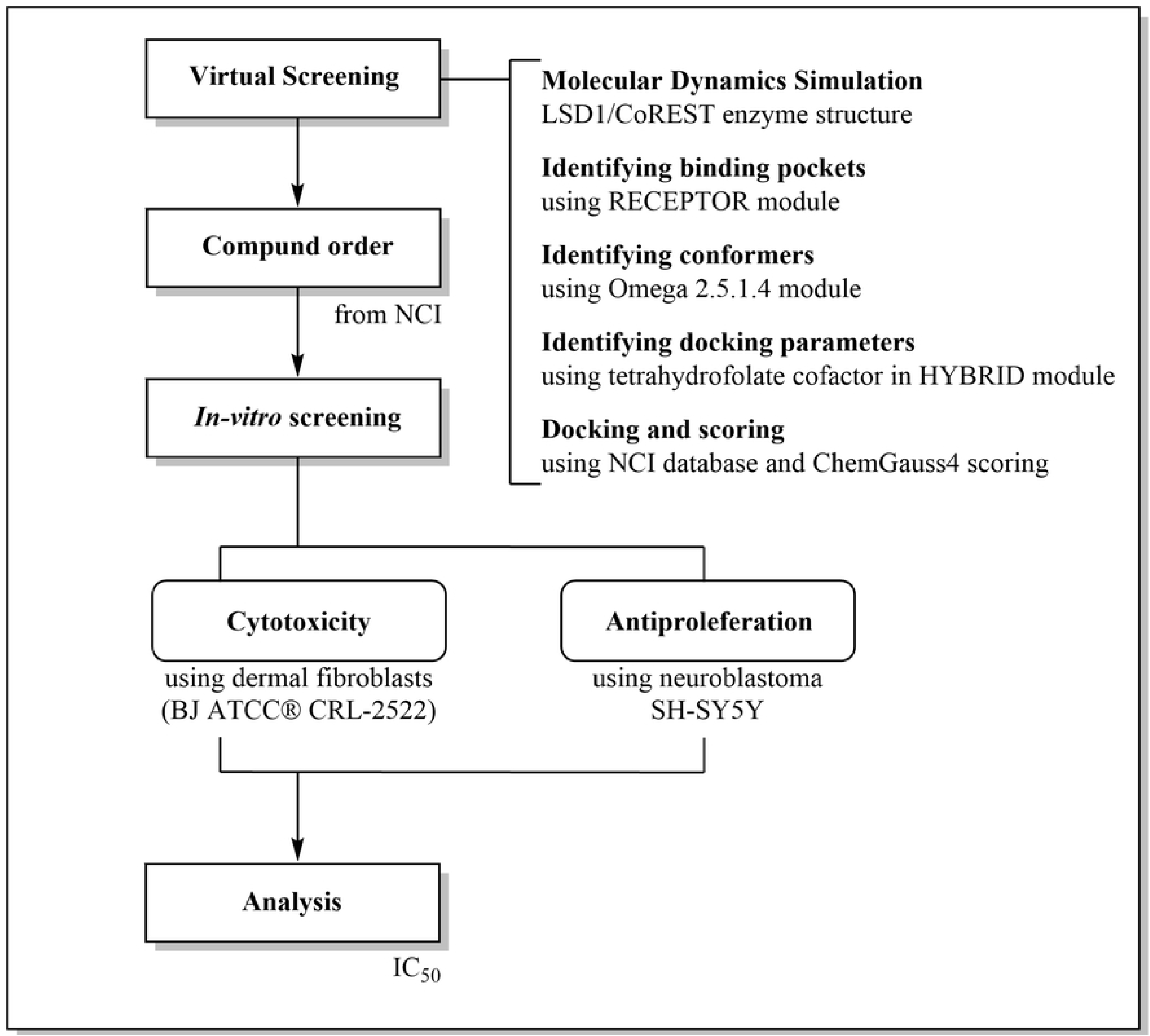
Study flowchart. LSD: Lysine-specific demethylase, NCI: National Cancer Institute, ATCC: American Type Culture Collection, IC50: Inhibitory concentration at which 50% of cells were killed.

## 3. Results

The exploration of the active center of LSD1/CoREST enzyme by long time molecular dynamics simulation (MD) revealed that its cofactor tetrahydrofolate (THF) binds to two binding sites. [36] The first site is located in the core of the binding center, while the second site located at the periphery near the CoREST domain. **Figure 2 A** and 2 **B** shows the binding interactions of THF to first and second sites respectively. According to these results, we used the energy-minimized structures resulted from the MD for the virtual screening of NCI database. **Figure 2 A** shows that THF mainly binds by hydrogen bonding in the first site. It binds to GLU559 by its amino group of the aromatic ring system, and to ILE356 and HIS564 backbone through one of its carboxylic acid moieties. Also, THF binds by hydrogen bonding to HIS564 side chain through its tail amide group. On the other hand, it forms hydrophobic interactions with VAL33 by aliphatic-aromatic forces and with HIS564 by π-π interactions. PHE538 also can form hydrophobic interactions with its aromatic ring system. **Figure 2 B** shows that THF bind by hydrogen bond to GLU387, ASN383, ASP556 amino acids through its aromatic ring system in the second binding site. The aromatic ring system also binds by ion-induced dipole to ARG837. THF binds by aromatic aliphatic interactions to LYS838 and THR561 side chains.

**Fig 2:**
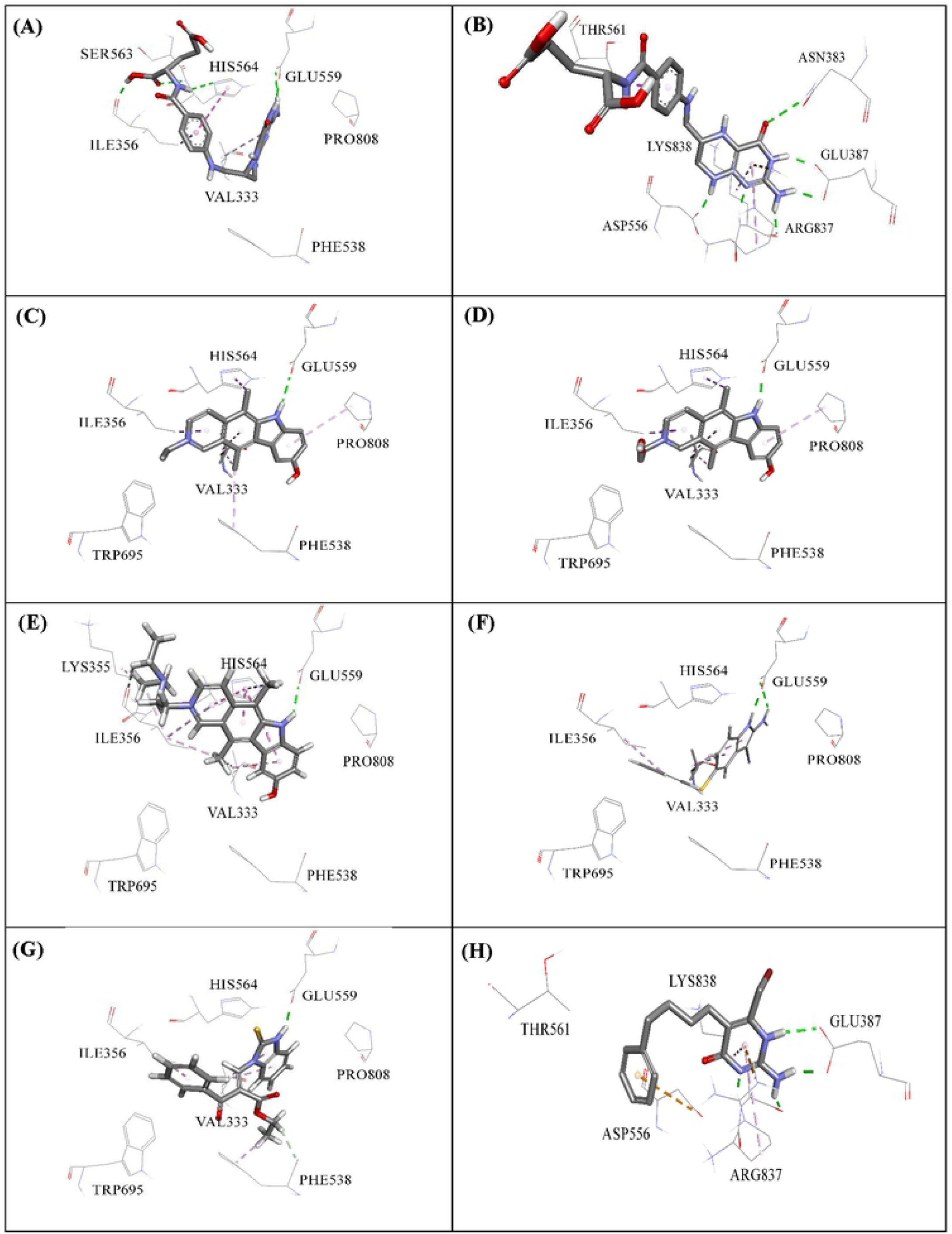
Binding of THF to the first site (**A**) and second site (**B**). Binding interactions of inhibitor compounds 1 (**C**), 2 (**D**), 3 (**E**), 4 (**F**) 5 (**G**), and 6 (**H**).

Based on the interactions represented by THF in both sites for experimental testing, *in-silico* screening generated 55 compounds. The obtained 55 compound were tested against the neuroblastoma cells at two concentrations (10 and 50 µM) as seen in **Table 1**. The compounds that showed more than 50 percentage of inhibition at both tested concentrations were considered as a potential ligands and their IC_50_ values against both neuroblastoma and normal fibroblast cell lines were determined. **Table 2** shows the structures of the best 5 tested compounds and their IC_50_ values against both cancer and normal cell lines.

**Table 1.** Percentage inhibition of neuroblastoma SH-SY5Y following 72 h exposure to 50μM and 10μM of all compounds.

| No# | NCS Code | % SH-SY5Y<br>50μM | % SH-SY5Y<br>10μM | No<br># | NCS Code | % SH-SY5Y<br>50μM | % SH-SY5Y<br>10μM |
| --- | --- | --- | --- | --- | --- | --- | --- |
| **1** | **NCI 311150** | **85.22 ± 0.004** | **82.78 ± 0.004** | 29 | NCI 379000 | 86.17 ± 0.003 | 15.08 ± 0.011 |
| **2** | **NCI 311151** | **86.63 ± 0.002** | **84.67 ± 0.001** | 30 | NCI 403185 | 41.03 ± 0.003 | 32.78 ± 0.003 |
| **3** | **NCI 311152** | **78.23 ± 0.057** | **74.22 ± 0.025** | 31 | NCI 522806 | 32.97 ± 0.002 | 27.75 ± 0.001 |
| **4** | **NCI 383336** | **58.93 ± 0.014** | **42.33 ± 0.010** | 32 | NCI 620256 | 42.03 ± 0.009 | 40.66 ± 0.000 |
| **5** | **NCI 655346** | **82.69 ± 0.005** | **41.30 ± 0.003** | 33 | NCI 655347 | 35.44 ± 0.002 | 30.40 ± 0.003 |
| **6** | **NCI 101194** | **51.31 ± 0.009** | **13.97 ± 0.003** | 34 | NCI 659751 | 74.08 ± 0.013 | 5.77 ± 0.001 |
| 7 | NCI 50930 | 23.73 ± 0.004 | 19.38 ± 0.002 | 35 | NCI 675224 | 32.51 ± 0.003 | 12.91 ± 0.002 |
| 8 | NCI 52363 | 20.97 ± 0.010 | 17.88 ± 0.007 | 36 | NCI 730608 | 76.37 ± 0.009 | 16.21 ± 0.000 |
| 9 | NCI 54112 | 31.91 ± 0.006 | 4.177 ± 0.001 | 37 | NCI 58561 | 28.66 ± 0.005 | 2.93 ± 0.003 |
| 10 | NCI 64862 | 36.68 ± 0.002 | 28.65 ± 0.0002 | 38 | NCI 62498 | 81.32 ± 0.002 | 6.97 ± 0.003 |
| 11 | NCI 79586 | 53.97 ± 0.001 | 23.98 ± 0.006 | 39 | NCI 67454 | 17.02 ± 0.003 | 12.88 ± 0.003 |
| 12 | NCI 88858 | 85.88 ± 0.001 | 33.41 ± 0.001 | 40 | NCI 83516 | 11.39 ± 0.007 | 2.02 ± 0.002 |
| 13 | NCI 88859 | 38.18 ± 0.001 | 24.39 ± 0.004 | 41 | NCI 107438 | 38.58 ± 0.002 | 35.08 ± 0.002 |
| 14 | NCI 90383 | 43.02 ± 0.001 | 16.12 ± 0.003 | 42 | NCI 130280 | 29.65 ± 0.003 | 26.89 ± 0.004 |
| 15 | NCI 92539 | 29.99 ± 0.008 | 21.30 ± 0.003 | 43 | NCI 141096 | 43.46 ± 0.003 | 37.57 ± 0.003 |
| 16 | NCI 92711 | 29.07 ± 0.004 | 13.45 ± 0.002 | 44 | NCI 210361 | 56.35 ± 0.004 | 32.32 ± 0.003 |
| 17 | NCI 108915 | 35.92 ± 0.001 | 6.27 ± 0.002 | **45** | NCI 210806 | 57.73 ± 0.007 | 30.57 ± 0.007 |
| 18 | NCI 112672 | 59.15 ± 0.002 | 14.37 ± 0.002 | 46 | NCI 211189 | 32.50 ± 0.005 | 1.56 ± 0.004 |
| 19 | NCI 115801 | 16.82 ± 0.004 | 13.05 ± 0.006 | 47 | NCI 211198 | 46.59 ± 0.004 | 28.91 ± 0.003 |
| 20 | NCI 121326 | 17.92 ± 0.006 | 14.86 ± 0.001 | 48 | NCI 211206 | 54.70 ± 0.005 | 4.60 ± 0.003 |
| 21 | NCI 131543 | 26.41 ± 0.006 | 18.87 ± 0.004 | 49 | NCI 211233 | 29.56 ± 0.003 | 0.55 ± 0.002 |
| **22** | NCI 131913 | 6.92 ± 0.002 | 1.72 ± 0.002 | 50 | NCI 280442 | 39.02 ± 0.01 | 31.78 ± 0.010 |
| **23** | NCI 136699 | 16.82 ± 0.004 | 13.91 ± 0.002 | 51 | NCI 291867 | 43.67 ± 0.037 | 32.95 ± 0.017 |
| 24 | NCI 142056 | 22.56 ± 0.006 | 13.36 ± 0.004 | 52 | NCI 336664 | 33.85 ± 0.030 | 32.95 ± 0.017 |
| 25 | NCI 179826 | 84.43 ± 0.007 | 21.62 ± 0.011 | 53 | NCI 605735 | 35.53 ± 0.015 | 29.07 ± 0.016 |
| **26** | NCI 196318 | 13.76 ± 0.006 | 14.31 ± 0.002 | 54 | NCI 615065 | 27.91 ± 0.013 | 21.58 ± 0.012 |
| 27 | NCI 251702 | 78.54 ± 0.007 | 21.30 ± 0.002 | 55 | NCI 698037 | 42.08 ± 0.013 | 21.27 ± 0.027 |
| 28 | NCI 346511 | 18.24 ± 0.002 | 9.67 ± 0.001 |  |  |  |  |

**Table 2:**
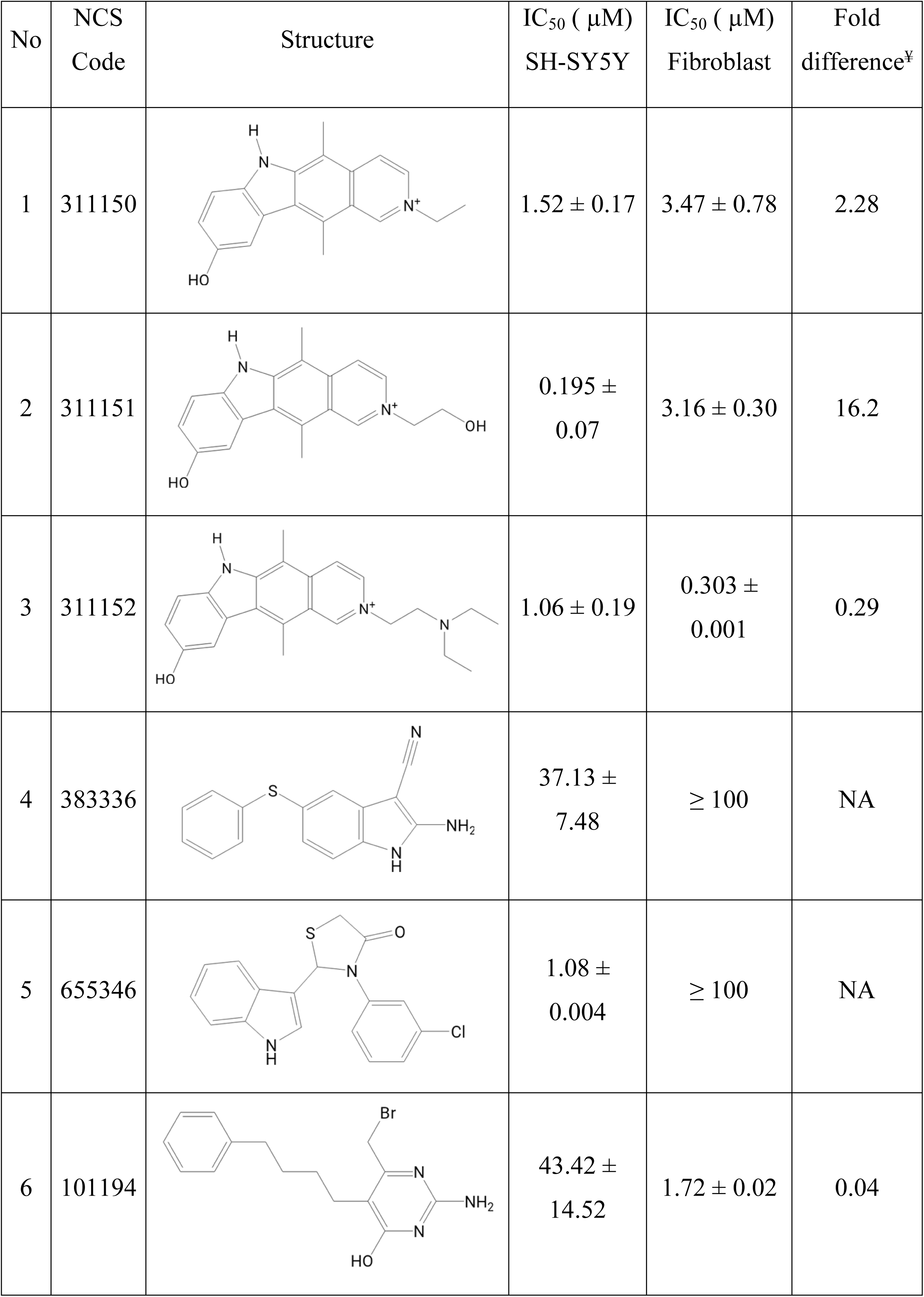
The IC_50_ values in µM for the tested most active compounds against neuroblastoma SH-SY5Y and normal fibroblast cell lines. ¥ fold difference represent the product of dividing IC_50_ in fibroblasts over the IC_50_ of SH-SY5Y. NA: not applicable.

| No | NCS Code | Structure | IC<sub>50</sub> ( μM)<br>SH-SY5Y | IC<sub>50</sub> ( μM)<br>Fibroblast | Fold<br>difference<sup>¥</sup> |
| --- | --- | --- | --- | --- | --- |
| 1 | 311150 |  | 1.52 ± 0.17 | 3.47 ± 0.78 | 2.28 |
| 2 | 311151 |  | 0.195 ± 0.07 | 3.16 ± 0.30 | 16.2 |
| 3 | 311152 |  | 1.06 ± 0.19 | 0.303 ± 0.001 | 0.29 |
| 4 | 383336 |  | 37.13 ± 7.48 | ≥ 100 | NA |
| 5 | 655346 |  | 1.08 ± 0.004 | ≥ 100 | NA |
| 6 | 101194 |  | 43.42 ± 14.52 | 1.72 ± 0.02 | 0.04 |

## 4. Discussion

Four compounds yielded and interestingly showed a relatively high selectivity index against the normal fibroblast cells. The selectivity index in the worst case was 2.5, which indicate that our compounds do exhibit a selectivity toward the LSD1 enzyme. Two of these compounds (compounds 3 and 6) were toxic on normal cells more than that on neuroblastoma cells and thus these compounds are not preferred for treatment of cancer. According to the protein atlas the expression level of the LSD1 in different types of the fibroblast cell lines range from 1028 to 26.2 while it is in the SH-SY5Y it is 42.2 [42].

Most of our compounds that gave promising activities upon testing against neuroblastoma SH-SY5Y belong to the virtual screening on the first site. **Figure 2** shows the binding of the most experimentally active compounds bound to the first (C, D, E, F and G) and second (H) binding sites. Comparing the interactions of the most active inhibitors to the binding site to that of the binding of THF to the first active site shows that they bind in the same manner. These compounds showed good experimental activities compared to other LSD1/CoREST reversible inhibitors. Also, binding of compound 6 to the active site of LSD1/CoREST second binding site is similar to that of THF binding according to the MD simulation.

Inspecting **Figure 2** shows that compounds 1 and 2 have interactions to PRO808, GLU559, HIS564, VAL333, ILE356 and PHE538. According to the docking pose and the interactions of both compounds, their activities should be similar. **Table 2** shows that the activity of compound 1 is 1.52µM while compound 2 has IC50 of 0.195µM. The difference between both compounds is only OH group at the side chain of compound 2, which does not form any kind of interactions to the active site. However, there is a huge difference in their activities, which is not explained by their interactions to the active site. Accordingly, we can explain the difference in their activities by their difference in delivery, where compound 2 has the extra OH group that decreased the intensity of the positively charged nitrogen atom, which enhances the delivery of this compound to the cytoplasm. Compounds 3 and 5 have activities comparable to compound 1. Compound 3 has structure similar to compounds 1 and 2 with an extended alkylamin at the side chain. Its activity is higher than that of compound 1, which may be assigned to the loss of the interaction with PRO808 and PHE538 amino acids. Its interaction pattern should predict a lower activity compound. The side chain has extra hydrophobic interaction to LYS355, which explains its higher activity. Although Compound 5 has activity very close to that of 3 compound, it has different structure from compounds 1, 2 and 3. This compound lacks the interactions with PRO808 and HIS564 amino acids and expected to have lower activity than compound 1, however it has a higher activity. Inspection the structure of this compound showed that it has a sulfur atom, which contributes to its lipophilicity and enhances its delivery. Compound 4 has lower activity than all abovementioned compounds, and it only interacts to GLU559, VAL333 and PHE538. The lower number of interactions this compound forms with the active site explains its low activity.

Compound 6 binds to the peripheral binding site of the THF co-enzyme. It binds with two hydrogen bonds with GLU387 through its imino and amino protons, which is the same pattern found in the binding of THF to the peripheral binding site (**Figure 2B**). This compound and THF bind to same amino acids through their heteroaromatic rings, where they can bind to ARG837 by ion-induced dipole interactions or aliphatic aromatic interactions with LYS838. Also, compound 6 bind to ASP556 through its benzene ring by ion-induced dipole interaction, which is the same interaction of THF but with its heteroaromatic ring. Compound 6 misses the interaction of THF to THR561 by aliphatic-aromatic interactions. The activity of this compound is not high as those bind to the first site, and we correlate that to the missed interactions to the binding site similar to THF. According to the models that were used in the virtual screening for this project, the solvation polar energy is higher for this binding site [33]. This shows that the peripheral site needs higher energy from the compound to bind due to its hydrophilic nature, which explains the low activity of compound 6.

Due to the potential of LSD1 as an anti-cancer target, several LSD1 inhibitors have been explored, and the majority of the very powerful LSD1 inhibitors have only recently been found [43, 44]. Peptides, trans-2-phenylcyclopropylamine derivatives, polyamines, and guanidine are among the inhibitors [45]. Most importantly, some KDM1 inhibitors have progressed to clinical trials for the treatment of leukemia and solid tumors, either alone or in combination with other treatments [46]. Such inhibitors are subcategories into five: MAO inactivators and their derivatives, natural products, peptide inhibitors, polyamine-based inhibitors, and metal-based inhibitors [40]. Out of those groups the peptide inhibitors were the most attractive with the SNAIL peptide-based molecule reaching an IC_50_ value of 0.28 μM and anti-proliferation effect for Hela cells [41]. In case of metal-based inhibitors, a rhodium (III) complex compounds was developed as the first metal-based inhibitor of LSD1 activity reported in the literature. This metal complex occupied the binding pocket of LSD1 for histone H3 recognition and thus blocked the LSD1-H3K4me2 interaction in human prostate cancer cells, leading to increasing amplification of p21, FOXA2, and BMP2 gene promoters with an IC_50_ of 0.04 ± 0.008 μM for LSD1 [47]. However, further work needs to be done to improve the bioavailability of the rhodium (III) complex in vivo. Comparing the compounds yielded from this work with what are available in the literature yield a good promise to our compounds in term of potency, selectivity, and safety.

## 5. Conclusions

The work represented in this study demonstrated that the molecular dynamics simulation can yield a selective inhibitor for LSD1/CoREST complex. Successful inhibitors may have a clinical application in treating cancers that overexpress the LSD1 enzyme such as the SH-SY5Y neuroblastoma that was tested in this work.

## Acknowledgements

The authors would like to thank the Abdul Hameed Shoman Foundation for funding this research, according to Agreement No. 05/2018 and the Deanship of Scientific Research at the University of Jordan. We would like also to acknowledge the Developmental Therapeutics Program, Division of Cancer Treatment and Diagnosis, National Cancer Institute (http://dtp.cancer.gov) for providing us with the NCI compounds.

## Availability of data and materials

All data are available upon reasonable request from the corresponding author.

## Authors’ contributions

### Ethics approval and consent to participate

This article does not contain any studies with human participants or animals performed by any of the authors.

### Patient consent for publication

Not applicable.

### Competing interests

The author(s) declare that they have no competing interests.

**Authors’ information (optional)** :

